# Epigenetic memory as a time integral over prior history of Polycomb phase separation

**DOI:** 10.1101/2020.08.19.254706

**Authors:** Jorine M. Eeftens, Manya Kapoor, Clifford P. Brangwynne

## Abstract

Structural organization of the genome into transcriptionally active euchromatin and silenced heterochromatin is essential for eukaryotic cell function. Heterochromatin is a more compact form of chromatin, and is associated with characteristic post-translational histone modifications and chromatin binding proteins. Phase-separation has recently been suggested as a mechanism for heterochromatin formation, through condensation of heterochromatin associated proteins. However, it is unclear how phase-separated condensates can contribute to stable and robust repression, particularly for heritable epigenetic changes. The Polycomb complex PRC1 is known to be key for heterochromatin formation, but the multitude of Polycomb proteins has hindered our understanding of their collective contribution to chromatin repression. Here, we take a quantitative live cell imaging approach to show that PRC1 proteins form multicomponent condensates through hetero-oligomerization. They preferentially seed at H3K27me3 marks, and subsequently write H2AK119Ub marks. Using optogenetics to nucleate local Polycomb condensates, we show that Polycomb phase separation can induce chromatin compaction, but phase separation is dispensable for maintenance of the compacted state. Our data are consistent with a model in which the time integral of historical Polycomb phase separation is progressively recorded in repressive histone marks, which subsequently drive chromatin compaction. These findings link the equilibrium thermodynamics of phase separation with the fundamentally non-equilibrium concept of epigenetic memory.

## INTRODUCTION

The genome encodes an organism’s heritable genetic information, but its differential expression in cells over time enables the variable phenotypic cellular behaviour that underlies biological function. The regulated expression of genes is known to be intimately linked to the structural organization of the genome, which plays a fundamentally important role in gene expression in all eukaryotic cells (1–3). In general, genomic sequences are packaged into two types of nuclear domains; euchromatin, an “open” state that allows for RNA transcription, and the more compacted heterochromatin, associated with inactive genes. The higher nucleosome density found in heterochromatin is thought to be inaccessible to transcriptional machinery, and refractory to chromatin remodelling required for transcription. Compaction is therefore widely accepted as a major hallmark of repressed chromatin, comprised of silenced genes which are not expressed (4, 5).

Heterochromatin can be further classified into two types, constitutive and facultative. Constitutive heterochromatin organizes repetitive sequences such as pericentromeric and subtelomeric regions into silent nuclear compartments, that are often located near the lamina or around nucleoli. In contrast, facultative heterochromatin consists of transcriptionally silent regions that can become active depending on the context (6). Constitutive heterochromatin is characterized by trimethylation of Histone H3 on lysine 9 (H3K9me3), and the presence of heterochromatin protein 1 (HP1), while facultative heterochromatin is characterized by trimethylation of Histone H3 on lysine 27 (H3K27me3) (6, 7). In addition to these distinct histone marks found in the two types of heterochromatin, facultative heterochromatin can be distinguished by the presence of a set of functionally important Polycomb proteins (8, 9).

The context-dependent silencing of facultative heterochromatin underscores the importance of understanding of how Polycomb proteins facilitate this process. Indeed, while Polycomb proteins are essential for development and cell differentiation, and also play a role in X-chromosome inactivation, cell cycle control, and maintenance of repression (10), much about how these multicomponent complexes facilitate compaction and silencing remains unclear. The canonical Polycomb Repressive Complex 1 (PRC1) consists of four core proteins: Cbx, PCGF, PHC, and RING, that each have numerous orthologs resulting in different compositions of PRC1 (9). The Cbx-subunit recognizes methylated Histone H3 on lysine 27 (H3K27me3), which is deposited there by the related Polycomb Repressive Complex 2 (PRC2). PRC1’s RING subunit, the E3 Ubiquitin ligase RNF2, subsequently deposits a second type of histone mark, by ubiquitinating H2AK119 (8, 11). PRC1 subunits have been shown to compact nucleosome arrays *in vitro* (12–14), although the biophysical mechanism driving this compaction is poorly understood.

Recent studies have provided support for the hypothesis that heterochromatin proteins may induce compaction through phase separation. Both HP1 and Polycomb proteins exhibit features characteristic of many proteins that drive phase separation, including oligomerization domains, intrinsically disordered regions, and substrate (chromatin) binding domains (15). Moreover, both are capable of undergoing liquid-liquid phase-separation, which through surface tension forces could directly drive chromatin into a more compact form (16–19). Phase separation of heterochromatin proteins could also function synergistically with chromatin, as several recent studies have shown that chromatin itself has an intrinsic ability to phase-separate and compartmentalize, in a manner which depends on particular histone marks (20, 21). Phase-separation could also potentially explain the mechanism behind heterochromatin spreading, a poorly understood phenomenon in which histone modifications defining the heterochromatin domain expand (22, 23). However, these chromatin marks are typically inherited by daughter cells after division (24–26), while most phase-separated condensates dissolve during mitosis (27). Indeed, the potential role of liquid-liquid phase separation in the reading and writing of histone marks, and whether phase separation is necessary for chromatin compaction, remains unclear.

Here, we show that through hetero-oligomerization, PRC1 Polycomb subunits drive the formation of multicomponent condensates, which can read and write repressive histone marks. Rather than PRC1 phase separation actively driving compaction, chromatin is instead compacted through the effect of subsequent post-translational modifications, particularly mediated by the ubiquitin ligase activity of RNF2, which is the central node of PRC1 subunit interactions. Thus, while phase separation arises from equilibrium thermodynamic driving forces which do not have memory, non-equilibrium reactions it facilitates can drive stable epigenetic changes, which serve to record the history of prior PRC1 phase separation.

## RESULTS

### Canonical PRC1 subunits form condensates

Polycomb proteins contain oligomerization domains, as well as other structural features (e.g. IDRs), characteristic of proteins driving phase separation (**Figure 1B,E,H,K**). The presence of oligomerization domains is particularly interesting, given recent studies highlighting the role of oligomerization in driving phase separation of condensates such as stress granules and nucleoli (28, 29). To understand the potential role of Polycomb protein oligomerization in promoting phase separation, we utilized the recently-developed Corelet system (30), a light-activated oligomerization platform (**Figure 1A**). Corelets consist of a 24-mer ferritin core, with each ferritin subunit bearing a photo-activatable iLID protein domain. Tagging a protein of interest with iLID’s binding partner sspB allows for its reversible, light-dependent oligomerization, which for some proteins can drive light-activated phase separation. Moreover, by varying the concentration ratio of core to sspB, the propensity of any protein to phase separate upon oligomerization can be precisely quantified, by mapping *in vivo* phase diagrams (30).

**Figure 1:**
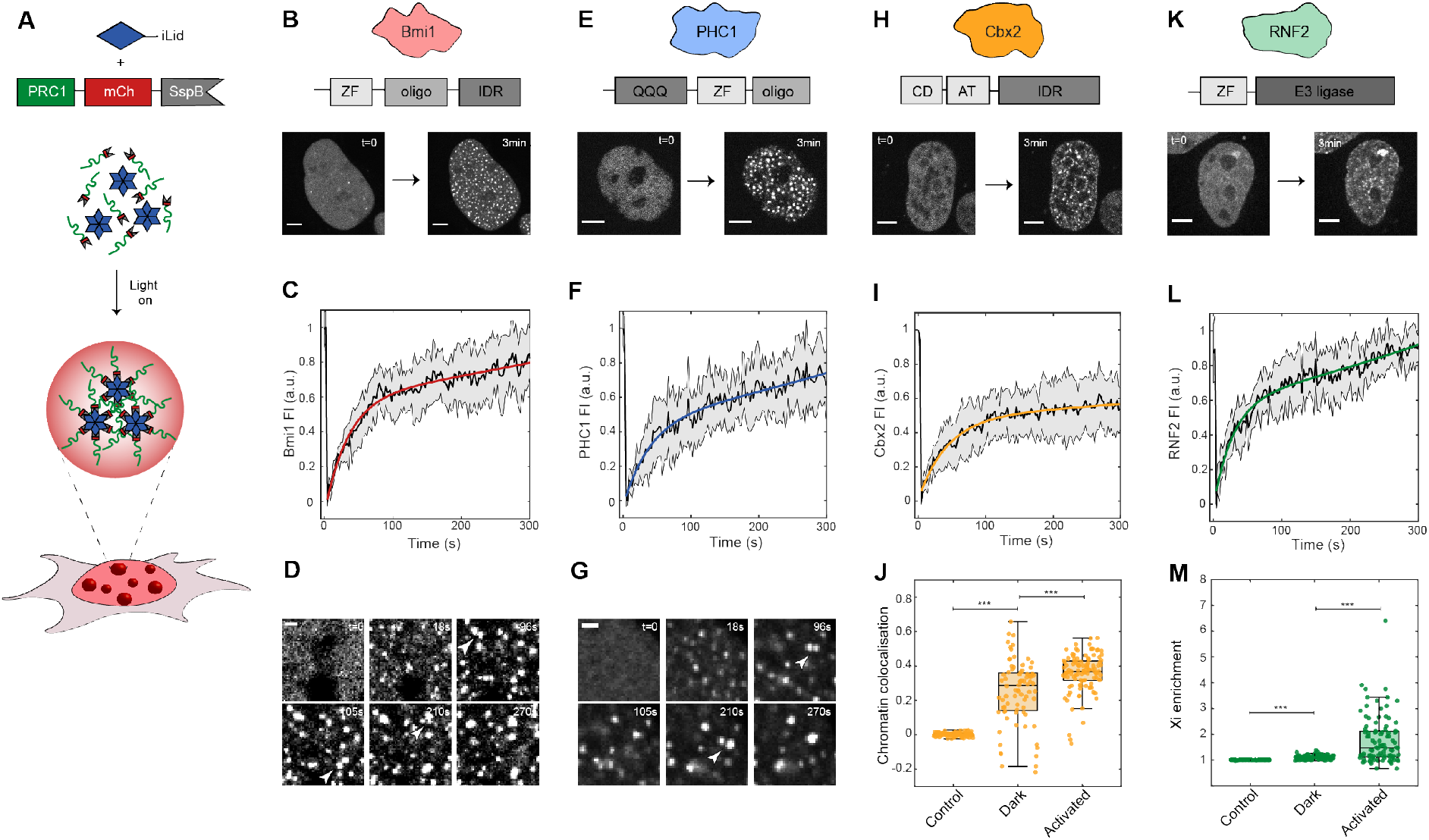
PRC1 subunit oligomerization drives phase separation. A: The Corelet system enables light-driven oligomerization to probe protein phase behaviour. B: Bmi1-mCh-sspB in HEK293 cells with NLS-Ferritin-Corelet before (t=0) and after blue light activation (3min). Scale bar is 5μm. C: FRAP recovery curve of Bmi1-mCh-sspB Corelets. Error bars represent standard deviation on at least 5 replicates. D: Bmi1 condensates fuse with one another. Arrows indicate condensates about to fuse. Scale bar is 1μm. E: PHC1-mCh-sspB in HEK293 cells with NLS-Ferritin-Corelet before (t=0) and after blue light activation (3min). Scale bar is 5μm. F: FRAP recovery curve of PHC1-mCh-sspB Corelets. G: PHC1 condensates fuse with one another. Arrows indicate condensates about to fuse. Scale bar is 1μm. H: Cbx2-mCh-sspB in HEK293 cells with NLS-Ferritin-Corelet before (t=0) and after blue light activation (3min). Scale bar is 5μm. I: FRAP recovery curve of Cbx2-mCh-sspB Corelets. J: Pearson correlation coefficient between chromatin (stained with Hoechst), and mCh-sspB (control), Cbx2-mCh-sspB (in the dark state), Cbx2-mCh-sspB (after activation). Cbx2-mCh-sspB preferentially localizes with chromatin. This colocalization is increased after activation. P-values<0.001. K: RNF2-mCh-sspB in HEK293 cells with NLS-Ferritin-Corelet before (t=0) and after blue light activation (3min). Scale bar is 5μm. L: FRAP recovery curve of RNF2-mCh-sspB Corelets. M: Partition coefficient of mCh-sspB (control), RNF2-mCh-sspB (in the dark state), RNF2-mCh-sspB (after activation) on the inactive X-chromosome. P-values<0.001.

We chose one ortholog of each of the canonical subunits (Bmi1/PCGF4, PHC1, Cbx2, RNF2/RING1B), and fused them to sspB, allowing us to interrogate the oligomerization-dependent phase behavior of each of these subunits in cultured human cells. We first examined Bmi1, which in addition to its N-terminal Zinc-finger (“ZF”) DNA binding domain, contains both an oligomerization domain and IDR (31) **(Figure 1B).** Before light activation, Bmi1 was mostly diffuse throughout the nucleus, with a few small areas of higher intensity, consistent with known endogenous localization patterns reported in literature **(Figure 1B)**(18, 19, 32–34). Upon light activation, many *de novo* condensates appeared, and the pre-existing puncta grew. The Bmi1-condensates recovered rapidly after photo bleaching, indicating a dynamic exchange of the majority of labelled Bmi1 molecules within ~2min, consistent with literature **(Figure 1C)**(33). While the Bmi1 puncta are close to the diffraction limit, they also appeared relatively round, and were frequently observed to undergo liquid-like coalescence with one another **(Figure 1D).** PHC1 is another PRC1 subunit with a native oligomerization domain, but in this case no significant predicted IDR. PHC1 exhibited behaviour similar to that of Bmi1: relatively diffuse prior to light activation, with *de novo* puncta after Corelet activation **(Figure 1E)**. PHC1 puncta also exhibited nearly complete FRAP recovery after several minutes and liquid-like fusion behaviour **(Figure 1FG)**. These data are consistent with Bmi1 and PHC1 having an inherent tendency to undergo liquid-liquid phase separation, at sufficient concentration and oligomerization.

Purified Cbx2 has previously been shown to undergo concentration-dependent phase separation *in vitro* (18, 19). However, it is unclear if the same is to be expected in the more complex intracellular environment, particularly given that Cbx2 contains a chromatin-binding chromo domain, and may thus be particularly subject to constraints from the presence of chromatin. Indeed, under Corelet activation, Cbx2 displayed behavior more complex than that of the clearly phase separated Bmi1 and PHC1 condensates. Prior to light activation, Cbx2 exhibited a variable intensity across the nucleus, consistent with previously reported endogenous localization (18, 19, 32) **(Figure 1H)**. Moreover, instead of forming distinct spherical droplets upon light activation, small chromatin-associated puncta formed, amplifying the pre-activated co-localisation pattern **(Figure 1J)**. These puncta exhibited FRAP recovery of only ~50% after >5min, indicating that these Cbx2 condensates contain a significant immobile fraction, presumably reflecting strong chromatin binding **(Figure 1I)**. RNF2-Corelets exhibited full FRAP recovery, but in this case also co-localizing with the inactive X-chromosome (Xi), with the colocalization enhanced upon light activation **(Figure 1K,L,M, Figure S1A)**.

Given that these PRC1 subunits exhibit a light-dependent amplification of their endogenous localization patterns and additional *de novo* puncta formation, we sought to further examine whether native intramolecular interactions are maintained. To test this, we expressed all subunits as GFP-fusion proteins, and assayed their recruitment into PRC1-Corelets, quantified by calculating the Pearson correlation coefficient (PCC) between the two fluorescent channels **(Figure 2, see Methods)**. For example, Bmi1-Corelet condensates strongly recruit PHC1-GFP, but not Cbx2-GFP **(Figure 2A)**. We found that condensates of each subunit could recruit like proteins, i.e. Bmi1-GFP localized into Bmi1-Corelet condensates, suggesting that each protein is capable of direct or indirect self-interaction **(Figure 2B,C,E,F, Figure S1B-E)**. Moreover, heterotypic interactions were relatively symmetric, e.g. Cbx2-Corelets recruited RNF2-GFP, and RNF2-Corelets recruited Cbx2-GFP **(Figure 2E,F)**, while PHC-Corelets do not recruit Cbx2-GFP, and Cbx2-Corelets do not recruit PHC1-GFP **(Figure 2C,E)**, underscoring the fidelity of the assay. Interestingly, while Bmi1, PHC1, and Cbx2 exhibit differential recruitment of the other subunits, RNF2 appears to recruit each with nearly equal apparent strength **(Figure 2F)**, suggesting it acts as a central node connecting the three other components, consistent with interaction studies (14, 32) **(Figure 2D)**. Thus, PRC1 Corelet constructs recapitulate key features of endogenous PRC1 complexes, and thus represent light-activatable, amplified versions of endogenous PRC1 condensates.

**Figure 2:**
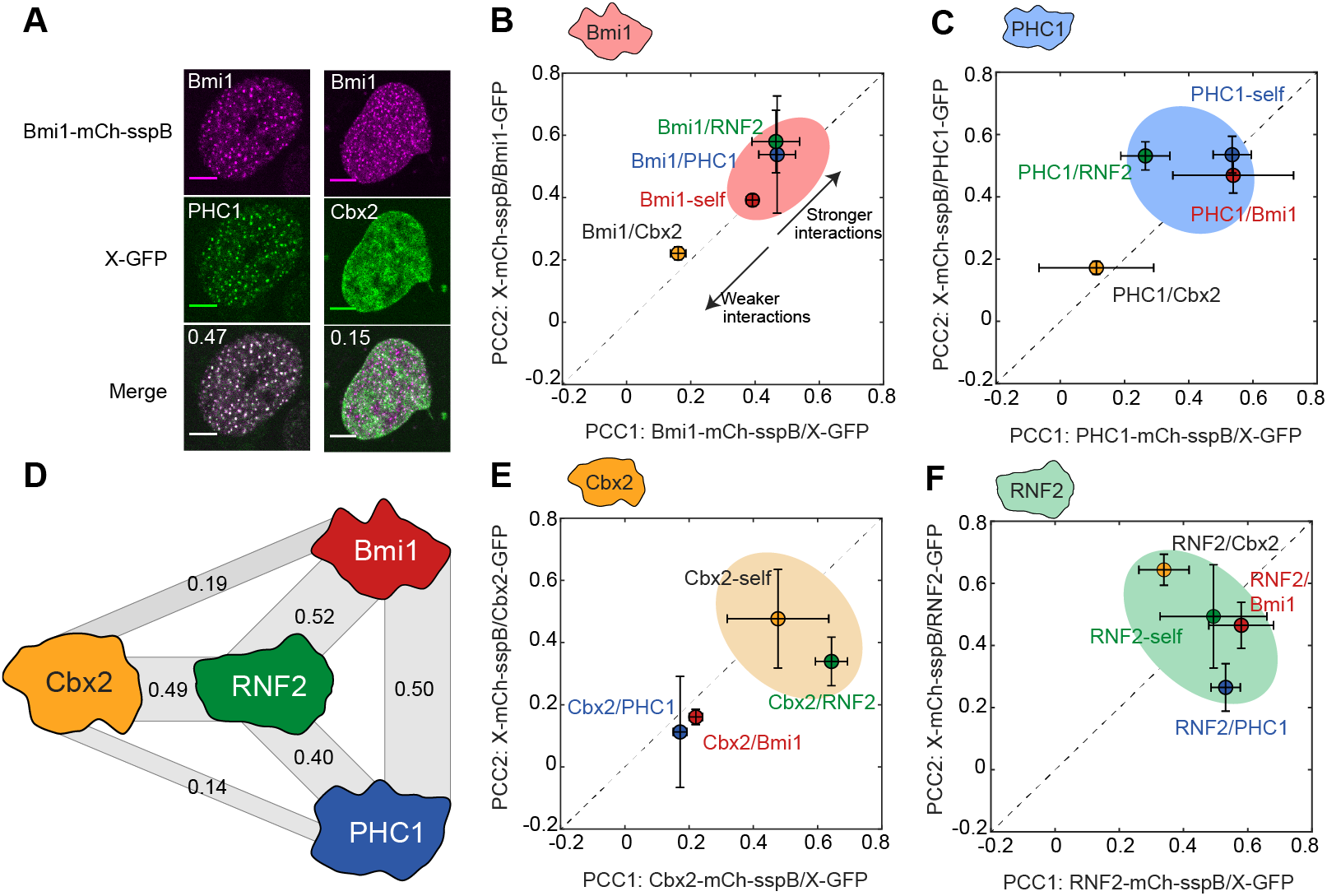
Engineered PRC1 condensates recruit endogenous partners. A: Examples of colocalization experiment. Bmi1-mCh-sspB Corelets recruit PHC1-GFP, but not Cbx2-GFP. Pearson correlation coefficient is shown in the upper left corner of the merged image. B: Recruitment of different PRC1 GFP fusion proteins into Bmi1-Corelets (PCC1) and Bmi1-GFP into different PRC1 Corelets (PCC2). Bmi1 shows self-interaction, and significant recruitment of PHC1 and RNF2. C: Recruitment of different PRC1 GFP fusion proteins into PHC1-Corelets (PCC1) and Cbx2-GFP into different PRC1 Corelets (PCC2). PHC1 shows self-interaction, and significant recruitment of Bmi1 and RNF2. D: Recruitment data suggests RNF2 is the central node in the PRC1 complex. Width of grey bars represents relative recruitment strength (average of the two PCC scores). E: Recruitment of different PRC1 GFP fusion proteins into Cbx2-Corelets (PCC1) and PHC1-GFP into different PRC1 Corelets (PCC2). Cbx2 shows self-interaction, and recruitment of RNF2. F: Recruitment of different PRC1 GFP fusion proteins into RNF2-Corelets (PCC1) and RNF2-GFP into different PRC1 Corelets (PCC2). RNF2 shows self-interaction, and significant recruitment of Bmi1, Cbx2, and PHC1.

### Hetero-oligomerization is responsible for condensate formation

We next sought to understand the relative contribution of native oligomerization domains, IDRs, and substrate-binding domains to PRC1 condensate formation. **(Figure 3A)**. We first focused on examining the contribution of the Bmi1 IDR to its phase behaviour, by truncating the protein to remove the IDR region (Bmi^ΔIDR^) **(Figure 3C)**. Interestingly, Bmi1^ΔIDR^-Corelets showed similar behaviour to the Bmi1^WT^. To precisely quantify any subtle difference, we mapped the intracellular phase-diagrams for both Bmi^WT^ and Bmi1^ΔIDR^. However, the phase-diagrams of the two are nearly identical, indicating that the tendency for intracellular phase-separation is not driven by the IDR **(Figure 3B,D)**. Moreover, Bmi1^ΔIDR^-Corelets could still recruit all other PRC1 subunits **(Figure 3E, Figure S2A)**, together indicating that the IDR region of Bmi1 contributes little to both PRC1 subunit recruitment and Polycomb condensate assembly. Interestingly, however, recruitment of full-length Bmi1-GFP was decreased, indicating Bmi1 exhibits weak interactions with itself, likely mediated by homotypic IDR-IDR affinity **(Figure 3E)**. By contrast, removal of Bmi1’s oligomerization domain (Bmi1^ΔOD^) completely abolished *de novo* Corelet condensate formation, with only slight growth of pre-existing puncta upon activation **(Figure 3FG)**.

**Figure 3:**
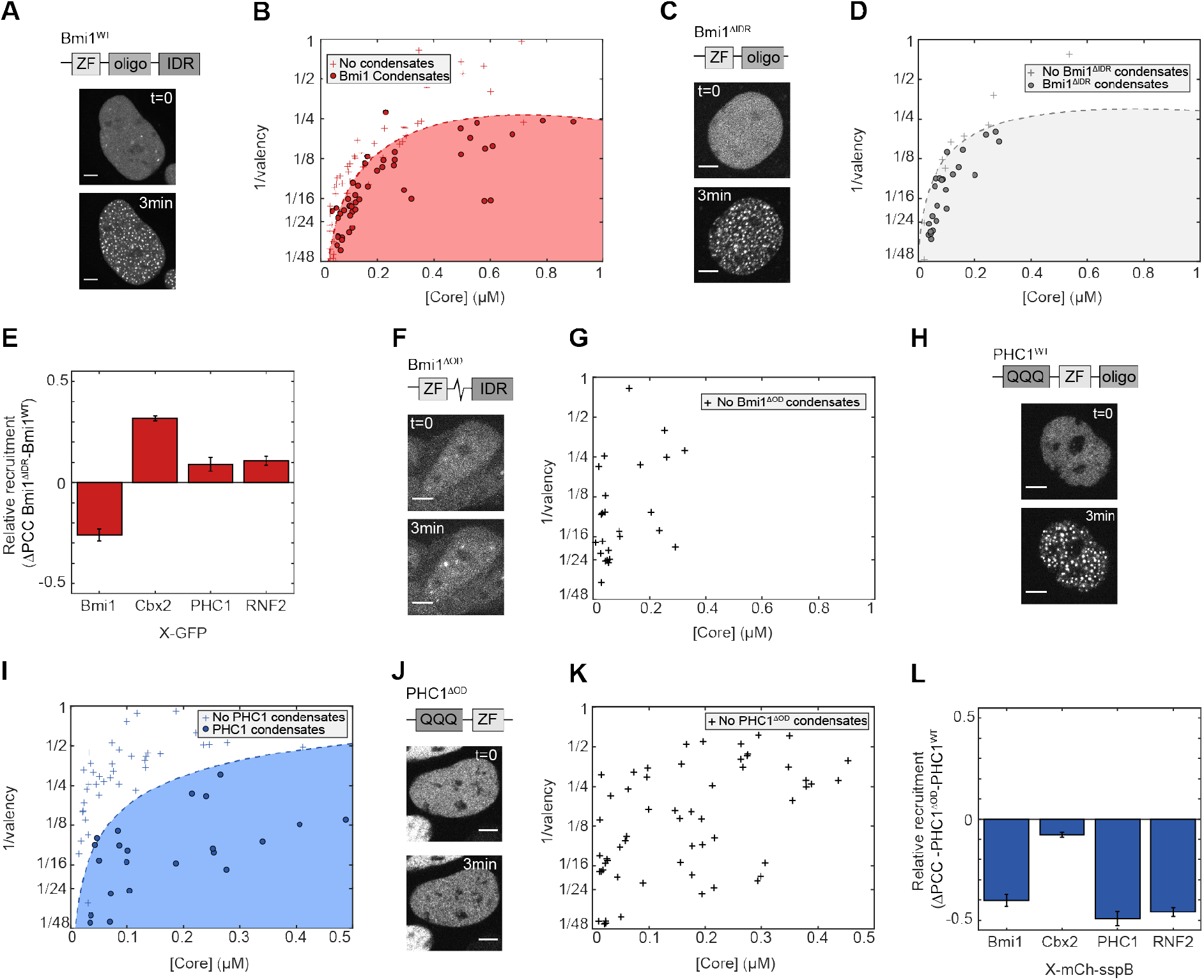
Hetero-oligomerization is responsible for condensate formation. A: Domain architecture of Bmi1^WT^ (top). Images show Bmi1^WT^-mCh-sspB in HEK293 cells with NLS-Ferritin-Corelet before (t=0) and after blue light activation (3min). Scale bar is 5μm. B: Phase-diagram of Bmi1^WT^. Filled red circles are phase-separated cells, red pluses are non-phase-separated cells. C: Domain architecture of Bmi1^ΔIDR^ (top). Images show Bmi1^ΔIDR^-mCh-sspB in HEK293 cells with NLS-Ferritin-Corelet before (t=0) and after blue light activation (3min). Scale bar is 5μm. D: Phase-diagram of Bmi1^ΔIDR^. Filled grey circles are phase-separated cells, grey pluses are non-phase-separated cells. E: Relative change in recruitment of PRC1-GFP fusions for Bmi1^WT^ and Bmi1^ΔIDR^. Self-interactions are decreased for Bmi1^ΔIDR^ (as indicated by negative ΔPCC), but recruitment of Cbx2-GFP is slightly increased (positive ΔPCC). Recruitment of PHC1 and RNF2 shows little change (ΔPCC close to zero). F: Domain architecture of Bmi1^ΔOD^ (top). Images show Bmi1^ΔOD^-mCh-sspB in HEK293 cells with NLS-Ferritin-Corelet before (t=0) and after blue light activation (3min). Scale bar is 5μm. G: Phase-diagram of Bmi1^ΔOD^, showing that Bmi1^ΔOD^ does not phase-separate in this regime. H: Domain architecture of PHC1^WT^ (top). Images show PHC1^WT^-mCh-sspB in HEK293 cells with NLS-Ferritin-Corelet before (t=0) and after blue light activation (3min). Scale bar is 5μm. I: Phase-diagram of PHC1^WT^. Filled blue circles are phase-separated cells and blue pluses are non-phase separated cells. J: Domain architecture of PHC1^ΔOD^ (top). Images show PHC1^ΔOD^-mCh-sspB in HEK293 cells with NLS-Ferritin-Corelet before (t=0) and after blue light activation (3min). Scale bar 5μm. K: Phase diagram for PHC1^ΔOD^, showing that PHC1^ΔOD^ does not phase-separate in this regime. L: Relative change in recruitment to PRC1-mCh-sspB fusions for PHC1^WT^ and PHC1^ΔOD^. Recruitment of all subunits is decreased for PHC1^ΔOD^ (negative ΔPCC). Neither PHC1^WT^ nor PHC1^ΔOD^ recruits Cbx2 (ΔPCC near zero).

It has been previously reported that oligomerization through the sterile alpha motif (SAM) domain of PHC1 plays a role in PRC1 clustering (35). First, we quantified the intracellular phase diagram for PHC1^WT^ **(Figure 3H,I)**. When we truncated PHC1 to remove the SAM-oligomerization domain, PHC1^ΔOD^-Corelets were also unable to form condensates (**Figure 3J,K)**. A comparison of the phase diagram of PHC1^ΔOD^ with PHC1^WT^ shows that PHC1^ΔOD^ is unable to form condensates in the concentration regime that the WT does **(compare Figure 3I with 3K)**. Thus, oligomerization by Corelets only, without the multiplicative effect of hetero-oligomerization with other PRC1 proteins, is insufficient for PHC1 phase separation. In addition, in contrast to full length PHC1, PHC1^ΔOD^-GFP was no longer recruited to any of the other PRC1 Corelet condensates **(Figure 3L, Figure S2B).** Thus, while IDRs appear largely dispensable, hetero-oligomerization domains are essential for multicomponent PRC1 condensate formation.

### PRC1 condensates recognize and write repressive histone marks

Cbx2 is known to read H3K27me3 marks, while RNF2 subsequently deposits a Ubiquitin mark (H2AK119Ub) **(Figure 4A)**(11). To examine whether this behaviour is recapitulated with our light-activated system, we first probed the interaction of Cbx2 condensates with H3K27me3. We fixed cells with Cbx2-Corelets before and after light activation and stained for histone marks with immunofluorescence. In non-activated cells, Cbx2 shows a variable intensity throughout the nucleus, with increased signal on denser chromatin regions **(Figure 4B, “OFF”)**. The H3K27me3 mark showed a more strongly punctate pattern throughout the nucleus, as well as a bright signal on the inactivated X-chromosomes. Upon light activation, there was no change in the H3K27me3 pattern, but Cbx2-condensates exhibit a clear preference for forming where these marks were, with a colocalization that increased with activation time (**Figure 4B)**. These cells contain unlabelled, endogenous Cbx2, and this colocalization could potentially reflect Cbx2 self-interactions, rather than H3K27me3 binding. To test this, we made a point mutation in Cbx2 that is known to prevent it from binding to the histone mark (Cbx2_F12A) (36). Although this mutated form still appeared to localize to the chromatin (presumably through the AT domain), this mutation reduced the overlap between Cbx2_F12A condensates and H3K27me3 to the level of a non-activated cell **(Figure 4B, S3A,B)**, consistent with Cbx2WT condensates colocalizing with H3K27me3 marks due to direct Cbx2 reading of these marks.

**Figure 4:**
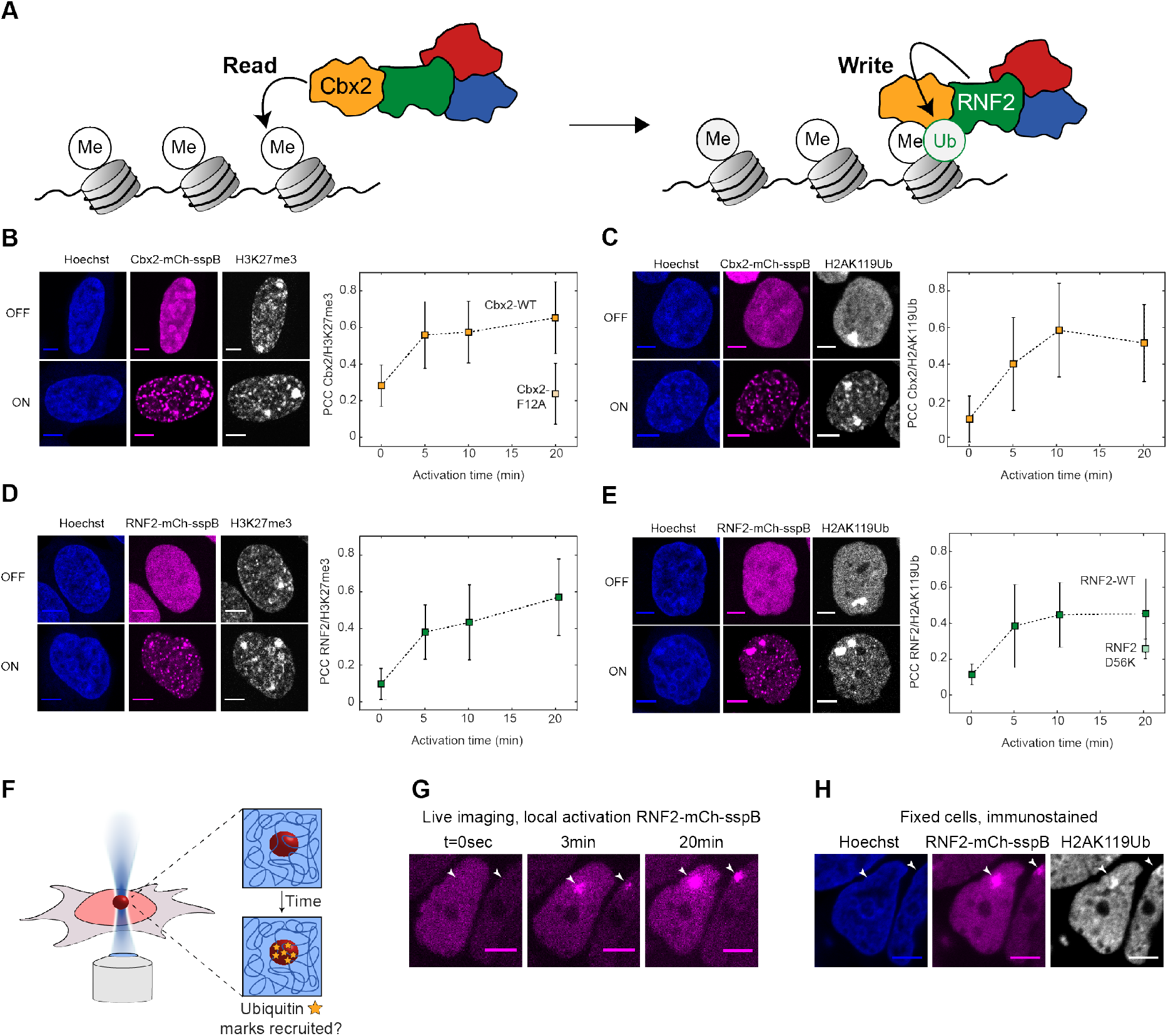
PRC1 condensates recognize and write repressive histone marks. A. The Cbx subunit of PRC1 recognizes the H3K27me3 mark, and the RING subunit writes the H2AK119Ub mark. B: Immunofluorescence on the H3K27me3 mark in fixed Cbx2-mCh-sspB Corelet cells, before (OFF) and after (ON) activation. Cbx2-mCh-sspB colocalizes with H3K27me3 marks upon light activation. The Cbx2_F12A-mCh-sspB mutant is unable to recognize the H3K27me3 mark. Scalebar is 5μm. Error bars represent standard deviation on at least 5 replicates. C: Immunofluorescence staining of the H2AK110Ub mark in fixed Cbx2-mCh-sspB Corelet cells, before (OFF) and after (ON) activation. Cbx2-mCh-sspB colocalizes with H2AK119Ub marks upon light activation. D: Immunofluorescence staining of the H3K27me3 mark in fixed RNF2-mCh-sspB Corelet cells, before (OFF) and after (ON) activation. RNF2-mCh-sspB colocalizes with H3K27me3 marks upon light activation. E: Immunofluorescence staining of the H2AK119Ub mark in fixed RNF2-mCh-sspB Corelet cells, before (OFF) and after (ON) activation. RNF2-mCh-sspB colocalizes with H2AK119Ub marks upon light activation. The RNF2_D56K-mCh-sspB mutant shows less colocalisation with the H2AK119Ub mark. F: Schematic of local activation experiment. Each cell is locally activated in a 1 micron^2^ area for 20 minutes, after which the cells are rapidly fixed. Fixed cells are immunostained with anti-H2AK119ub antibody, and stained with Hoechst. G: Local activation of RNF2-mCh-sspB. Arrows indicate region of interest. H: Fixed cells after local activation of RNF2-mCh-sspB, stained with Hoechst and anti-H2AK119Ub immunofluorescence.

We next sought to examine the relationship between the Cbx2-recruiting H3K27me3 marks, and H2AK119Ub marks. Cbx2 is not directly responsible for writing the H2AK119Ub mark, but since Cbx2 Corelet condensates strongly recruit the H2AK119Ub writer RNF2 **(Figure 2E, Figure S1D)**, we reasoned that these condensates could potentially give rise to H2AK119Ub marks. Before activation, H2AK119Ub is only prominently present on inactivated X-chromosomes **(Figure 4C, ‘OFF’)**. However, upon activation and condensation of Cbx2 condensates, H2AK119Ub marks begin appearing over time, at the same location where Cbx2 condensates form **(Figure 4C).** Thus, Cbx2 condensates are recruited by H3K27me3 marks, and can subsequently facilitate the writing of ubiquitination marks onto chromatin within the condensate.

To further examine the causal relationship between H3K27me3 and H2AK119Ub marks, we examined their localization with respect to RNF2 Corelets. Before activation, RNF2 was homogenously distributed, while the H3K27me3 mark showed the familiar punctate pattern, with increased signal on the inactive X-chromosomes **(Figure 4D, ‘OFF’)**. Upon light activation, RNF2 condensates began forming at the H3K27me3 marks, similarly to Cbx2 **(Figure 4D)**. Moreover, as with Cbx2 condensates, over time H2AK119Ub marks appear where RNF2-condensates formed **(Figure 4E)**. Consistent with RNF2 directly writing this ubiquitin mark, Corelets formed with an enzymatically dead mutant, RNF2_D56K, showed a slightly reduced colocalisation with the H2AK119Ub mark, although the effect was small, likely due to RNF2_D56K retaining an ability to recruit endogenous WT RNF2, and other PRC RING components **(Figure 4E, Figure S3C,D)**(37, 38). Thus, Cbx2 and RNF2 collaborate to read and write the histone code.

We next sought to determine whether the H2AK119Ub marks could be written at arbitrary genomic locations. A powerful feature of the light-activatable Corelet system is that we can locally activate within subregions of the nucleus, by focusing the 488 nm laser on a small region of interest **(Figure 4F)**. We activated a single 1-micron^2^ area in each nucleus for 20 minutes, fixed the cells and immunostained for H2AK119Ub **(Figure 4G,H).** Remarkably, the H2AK119Ub mark appeared specifically in the locally activated RNF2-condensates (**Figure 4H, Figure S3E).** We also confirmed that these bright H2AK119Ub marks do not simply reflect inadvertent activation on inactive X-chromosomes **(Figure S3E)**. Thus, inducing local phase separation of PRC1 condensates at any chosen genomic location results in writing of repressive histone marks there.

### PRC1 condensates lead to chromatin compaction

We next sought to utilize our light-inducible PRC1 Corelets to examine the relationship between phase-separation, epigenetic mark reading and writing, and chromatin compaction. **(Fig. 5A).** We investigated whether Cbx2-Corelets could compact the DNA by visualizing the chromatin distribution with an miRFP-tagged Histone2B (H2B-miRFP). Both Cbx2-mCh-sspB and H2B-miRFP appear fairly uniform in the dark state **(Figure 5B, t=0sec)**. Upon activation, Cbx2-condensates form, with a variance in the signal that rapidly increases, stabilizing over 5-10min **(Figure 5C)**. By contrast, upon activation, the variance in the H2B signal increases in a roughly linear fashion, continuing to increase even after 15min, when the Cbx2 condensates have fully formed **(Figure 5D)**. Moreover, when Cbx2 activation is terminated, the Cbx2-condensates rapidly disappear within several minutes **(Figure 5C),** but the compaction remains, even after another 20-30min **(Figure 5D).** Thus, while condensates steadily promote chromatin compaction, their presence is not required for maintenance of the compacted state. Consistent with this picture, the H2B variance appears to be an integral of the Cbx2 variance, such that compaction effectively sums over the prior history of PRC1 phase separation **(Figure 5E)**. Fixed cells stained with Hoechst at increasing time points show the same trend, indicating this is not an artefact of tagged H2B (**Figure S4AB)**. The H3K27me3-binding deficient Cbx2_F12A-condensates show the same behaviour as the WT upon activation, with rapidly increasing variance that levels off, and rapidly decreases upon deactivation **(Figure S4EF)**. But in contrast to the Cbx2-WT, the mutant is unable to compact chromatin, indicating that interaction with nucleosomes is crucial for this effect (**Figure S4G)**. Interestingly, however, RNF2-Corelets only exhibit a modest triggering of compaction **(Figure S4CD)**. We hypothesize this is due to the limited size of RNF2 condensates.

**Figure 5:**
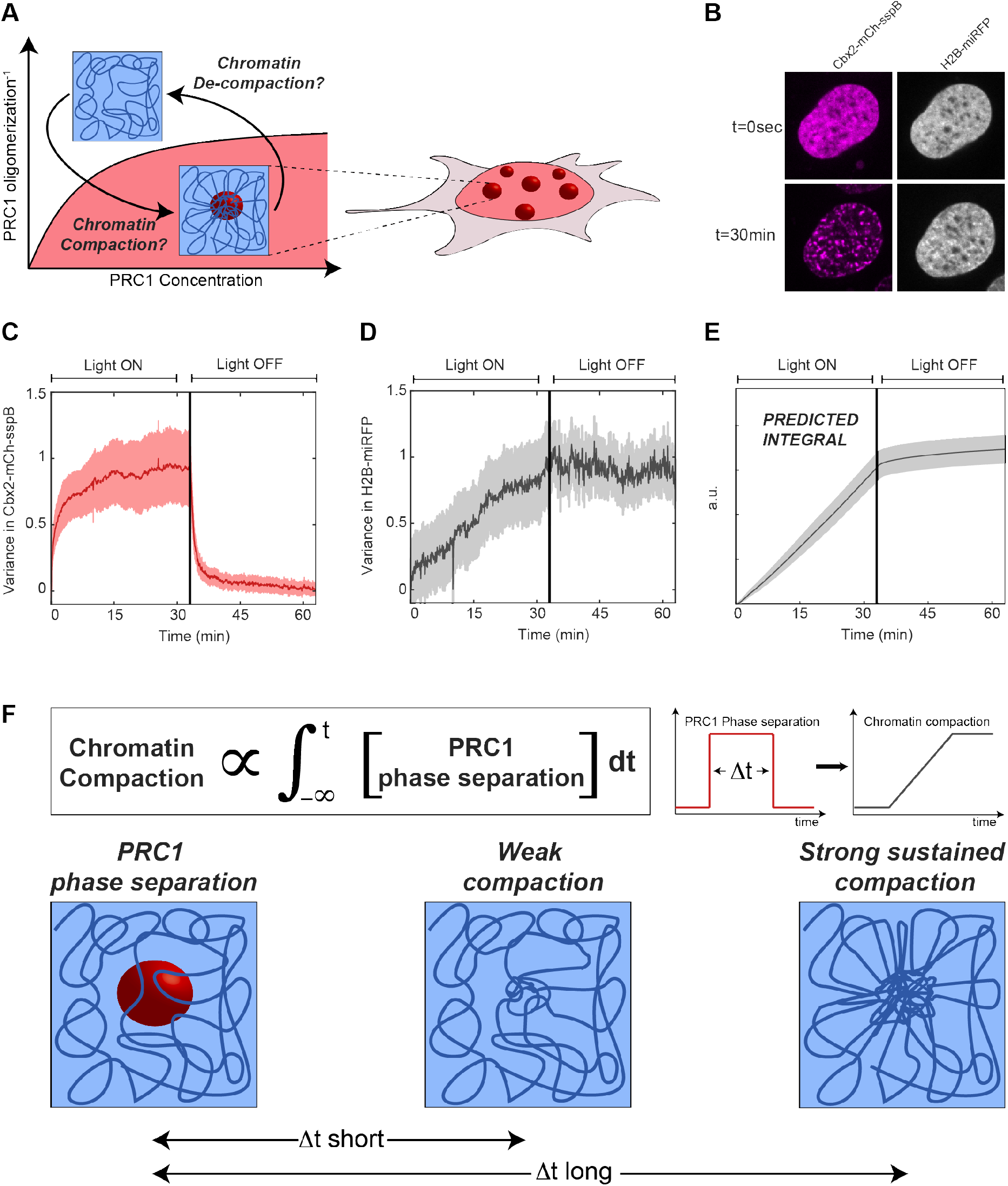
PRC1 condensates lead to chromatin compaction. A: Schematic of compaction experiment. After inducing condensate formation, chromatin compaction is examined, and tested to determine if compaction remains after deactivation. B: Cbx2-Corelet cell with H2B-miRFP before (t=0) and after activation (30min). C: The normalized variance in the Cbx2 signal increases rapidly as condensates form, and decreases rapidly upon deactivation, as condensates dissolve. D: The normalized variance in the H2B-miRFP signal increases gradually over time, and the high variance remains after deactivation, indicating sustained compaction. E: Integral of the Cbx2 variance, showing that compaction sums over the prior history of Cbx2 phase separation F: Mathematical relationship between polycomb phase separation, and degree of chromatin compaction. Long durations of prior PRC1 phase separation can lead to strong sustained chromatin compaction, even in the absence of sustained PRC1 phase separation.

## DISCUSSION

Here we have used quantitative live cell imaging approaches probe the interplay between phase separation, compaction, and post-translational modifications in the process of heterochromatin formation in living cells. We developed light-activated, multicomponent PRC1 condensates, which allow mapping of subunit recruitment and collective propensity for phase separation. Consistent with their ability to recapitulate key features of endogenous PRC1, we showed that these engineered condensates can directly recognize and write histone marks associated with facultative heterochromatin. Using the powerful spatiotemporal control of this system, we demonstrate that PRC1 condensation can induce chromatin compaction, but sustained phase separation is dispensable for maintenance of the compacted state.

Our approach enabled the interrogation of the role of each of the four core subunits of PRC1, and specific domains of these proteins, in driving phase separation. Strikingly, while Bmi1 robustly phase separates into liquid-like condensates upon oligomerization, its IDR did not have a significant impact on condensate formation. PHC1 shows similar phase separation behaviour, while it lacks a significant IDR. This is consistent with recent work showing that while weak interactions between IDRs can drive phase-separation in certain systems (39–44), a more complicated picture is emerging for multicomponent systems in living cells (28, 45).

Our light activated Corelet system confirmed the previously reported interactions between PRC1 components (46, 47), while highlighting a heterotypic network of weak affinities among subunits, that drive multicomponent PRC1 condensates formation. This is a more complicated picture than a previously proposed client-scaffold model, in which Cbx2 would function as the liquid scaffold, with the rest of the PRC1 components as clients (48). While each of the PRC1 subunits has an intrinsic capability to form condensates, Cbx2 is actually the least dynamic in nature. Instead, Cbx2 may acts as a localizing “seed” on the chromatin, and through its interactions with the other phase separation-prone PRC1 subunits, particularly Bmi1 and PHC1, locally amplifies valence to drive formation of a larger condensate. This local valence is further amplified through oligomerization domains on Bmi1 and PHC1, which together with RNF2 appear to serve as valence-amplifying “nodes” (28, 35, 49, 50).

As the catalytic subunit, the E3 ubiquitin ligase RNF2 thus plays a unique role, both as a structural component linking Bmi1 and PHC1 to the “reader” Cbx2, and also functioning as the “writer”, to translate PRC1 condensate formation into localized repressive Ubiquitin marks.

Polycomb condensates have previously been proposed to function by bringing distant chromatin loci together (38, 51–54). This picture distinguishes the catalytic activity (Cbx2, RNF2) of PRC1 from its potential role in chromatin remodelling, which could rely on strong oligomerizing of Bmi1 and PHC1 (35, 51, 53). If two genomic loci are bound by a Polycomb condensate, surface-tension mediated pulling of the two loci could bring the loci together, while excluding parts of chromatin not bound by Polycomb proteins (44, 55). Thus, PRC1 condensates could potentially serve to directly compact the genome, by pulling genomic elements together into dense clusters.

Despite the attractive simplicity of this PRC1-condensation/compaction picture, our data are inconsistent with PRC1 condensates directly compacting chromatin. Indeed, upon induction, chromatin continues steadily increasing in compaction, even after the PRC1 condensates have stabilized. The compaction level appears to reflect a time integral over the condensates. This summation of prior exposure to phase separated PRC1 is consistent with PRC1 condensates serving as reaction crucibles. These findings echo those in a recent study that showed that H2B ubiquitination enzymes work together to form a “reaction-chamber condensate”, possibly in a similar manner to PRC1 (56). Consistent with this alternative picture, even after our induced condensate dissolves, the modified chromatin remains in its compacted form **(Figure 5D),** underscoring how compaction cannot be the result of physical force directly imparted by the condensate. Instead, we show that induced condensates lead to an increase in repressive histone marks. These repressive histone marks (both H3K27me3 and H2AK119Ub), rather than the condensates themselves, subsequently drive compaction of chromatin.

One possible explanation for how modified chromatin becomes compacted, is chromatin’s intrinsic ability to compartmentalize itself, a possibility that has gained support in several recent studies (20, 21). We envision that as long as the condensates are present, the dynamic exchange of proteins within them leads to ubiquitination of the interwoven chromatin, which subsequently compacts itself. Moreover, as the chromatin compacts, new histones move into the condensate, and are subsequently modified to further drive compaction. This picture would explain the observed linear increase in compaction even after the condensates growth saturates, and maintenance of the compacted state after the condensates are dissolved, a result echoed in recent experiments on the role of HP1 in constitutive heterochromatinization (57).

Phase separation in biology has gained considerable attention in recent years, including as a potential mechanism for epigenetic changes. However, as an equilibrium thermodynamic framework, phase separation alone cannot explain heritable epigenetic changes, which must be robustly maintained through replication and cell division. Our results reconcile this key discrepancy, by showing how the equilibrium process of liquid-liquid phase separation can lead to long-lasting non-equilibrium effects, underscoring the complex interplay between the physicochemical driving force of phase separation and the reading and writing of epigenetic information.

## ACKNOWLEDGEMENTS

We thank Mike Levine, as well as members of the Brangwynne lab for discussions and comments on the manuscript, including Dan Bracha, Josh Riback, and Amy Strom. This work was supported by the Howard Hughes Medical Institute, and grants from the NIH 4D Nucleome Program (U01 DA040601). J.M.E. is supported by a Rubicon Grant (NWO).

## AUTHOR CONTRIBUTIONS

Conceptualization: J.M.E. and C.P.B. Software: J.M.E. Formal analysis: J.M.E. Investigation: J.M.E. and M.K. Writing-Original Draft: J.M.E. and C.P.B. Writing – Review and Editing: all authors. Supervision: C.P.B.

---

**Figure S1:**
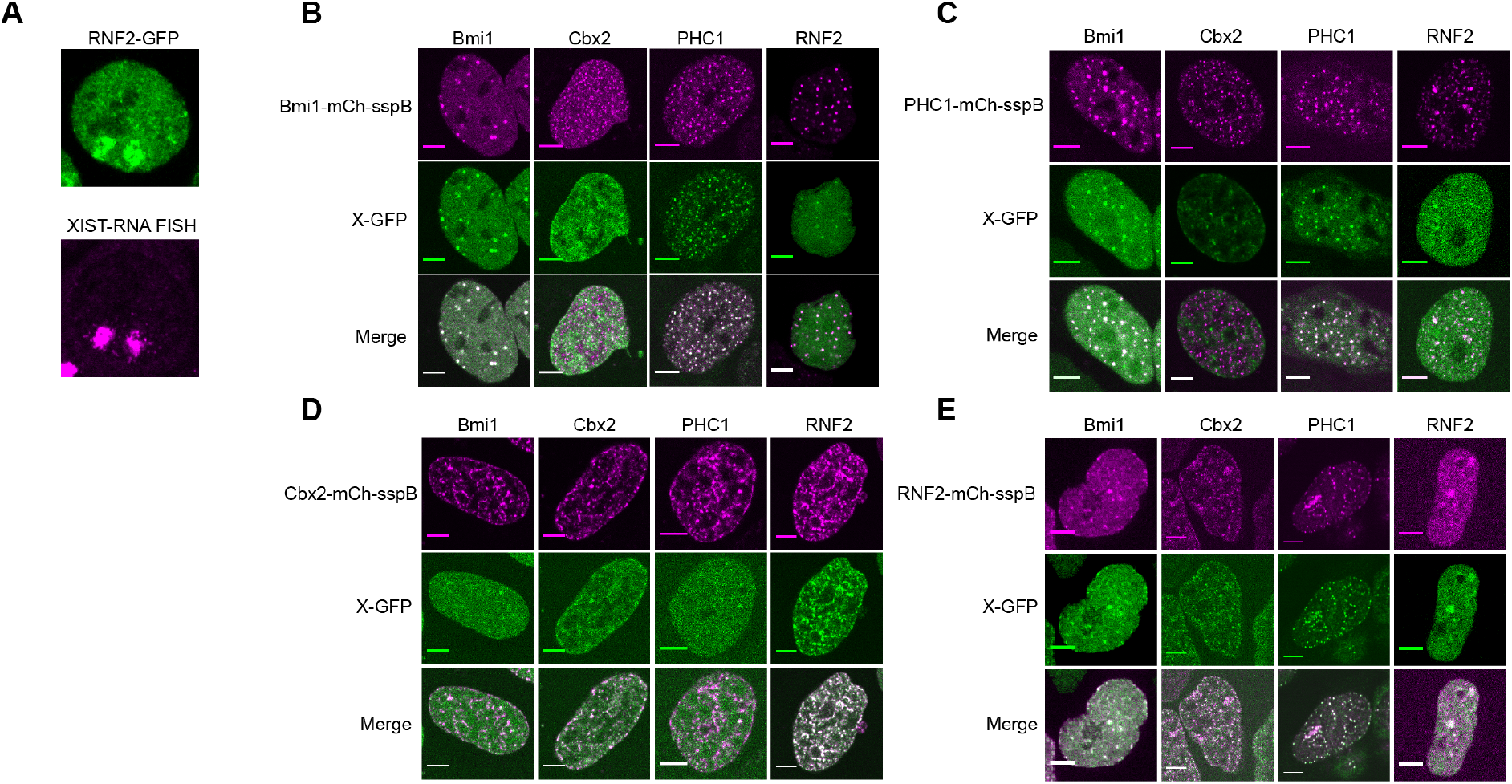
A. RNA-FISH on XIST shows that RNF2 partitions onto the inactive X-chromosomes. B. Recruitment of PRC1 subunit GFP-fusions to Bmi1-mCh-sspB Corelets. Cells were activated for 3 minutes. Scalebar is 5μm. C. Recruitment of PRC1 subunit GFP-fusions to PHC1-mCh-sspB Corelets. D. Recruitment of PRC1 subunit GFP-fusions to Cbx2-mCh-sspB Corelets. E. Recruitment of PRC1 subunit GFP-fusions to RNF2-mCh-sspB Corelets.

**Figure S2:**
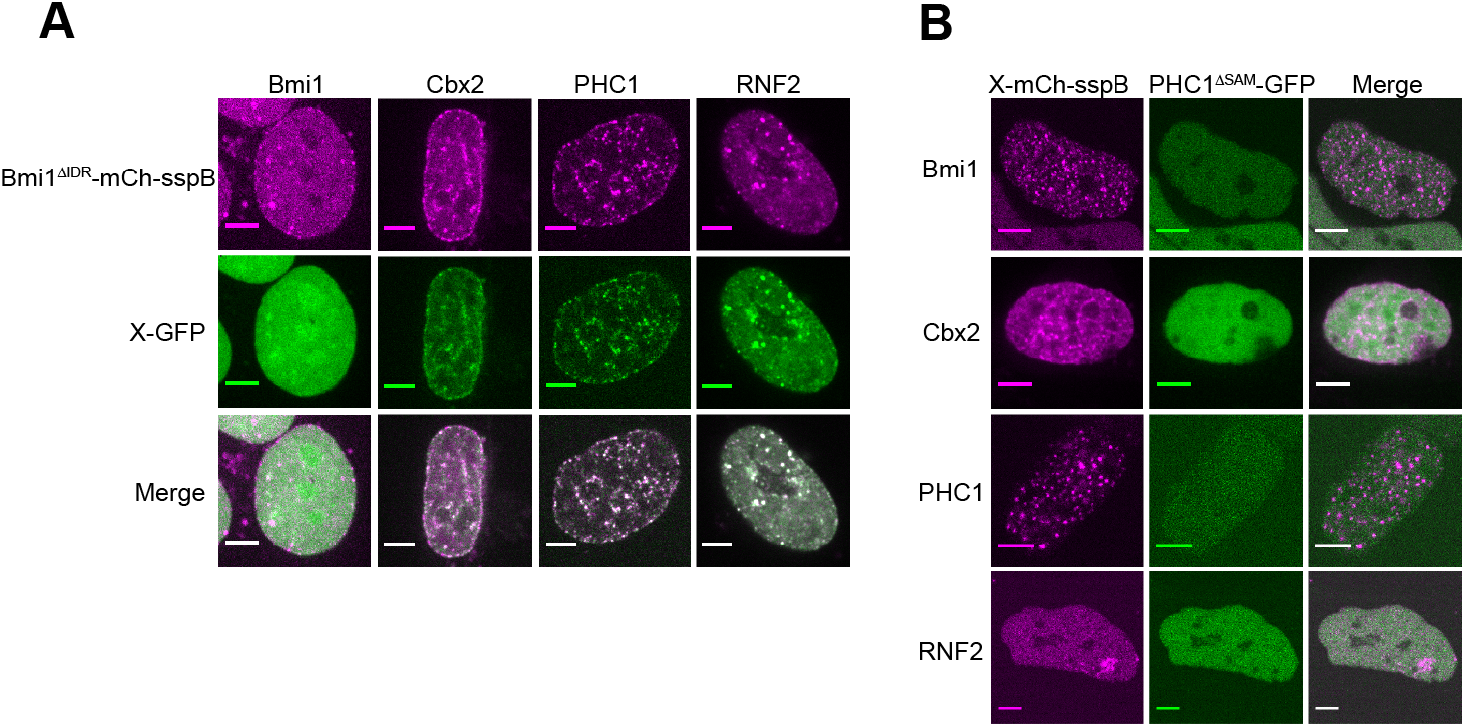
A. Recruitment of PRC1 subunits to Bmi1^ΔIDR^-mCh-sspB Corelets. Cells were activated for 3 minutes. Scalebar is 5μm. B. Recruitment of PHC1^ΔOD^-GFP to PRC1-mCh-sspB Corelets.

**Figure S3:**
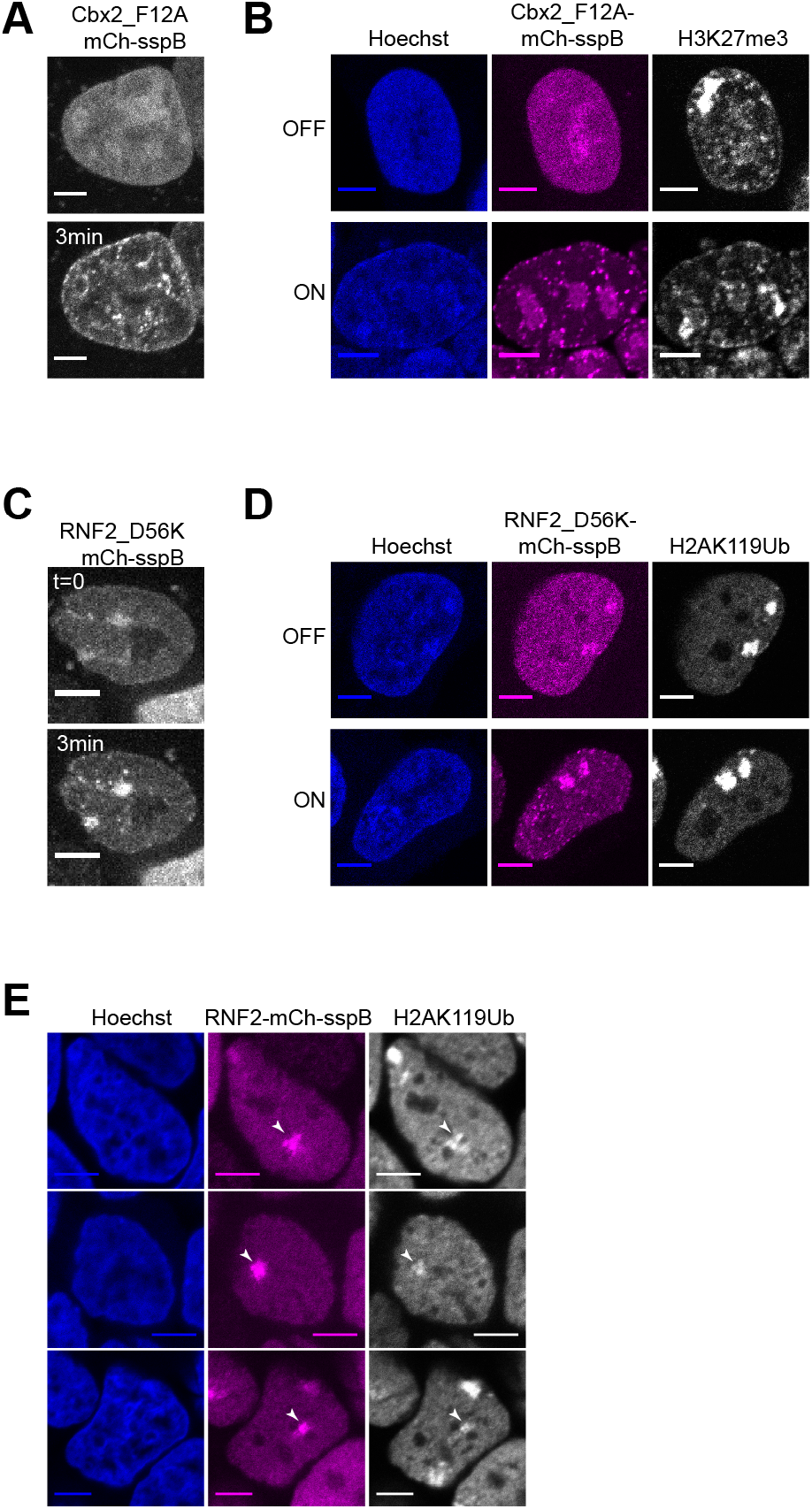
A. Cbx2^F12A^-mCh-sspB (point mutation abolishes the ability to recognize the H3K27me3 histone mark) in HEK293 cells with NLS-Ferritin-Corelet before (t=0) and after blue light activation (3min). Scale bar is 5μm. B. Immunofluorescence on the H3K27me3 mark in fixed Cbx2^F12A^-mCh-sspB Corelet cells, before (OFF) and after (ON) activation. Cbx2^F12A^ does show increased colocalization with H3K27me3 marks. C. RNF2^D56K^-mCh-sspB (point mutation that abolishes the ability to write the H2AK119Ub mark) in HEK293 cells with NLS-Ferritin-Corelet before (t=0) and after blue light activation (3 min). D. Immunofluorescence on the H2AK119Ub mark in fixed RNF2^D56K^-mCh-sspB Corelet cells, before (OFF) and after (ON) activation. RNF2^D56K^ moderately localizes with H2AK119Ub marks. E. Three examples of the H2AK119Ub mark appearing where RNF2 is locally activated. In the bottom example, two inactive X-chromosomes are clearly distinguishable in addition to the activated spot.

**Figure S4:**
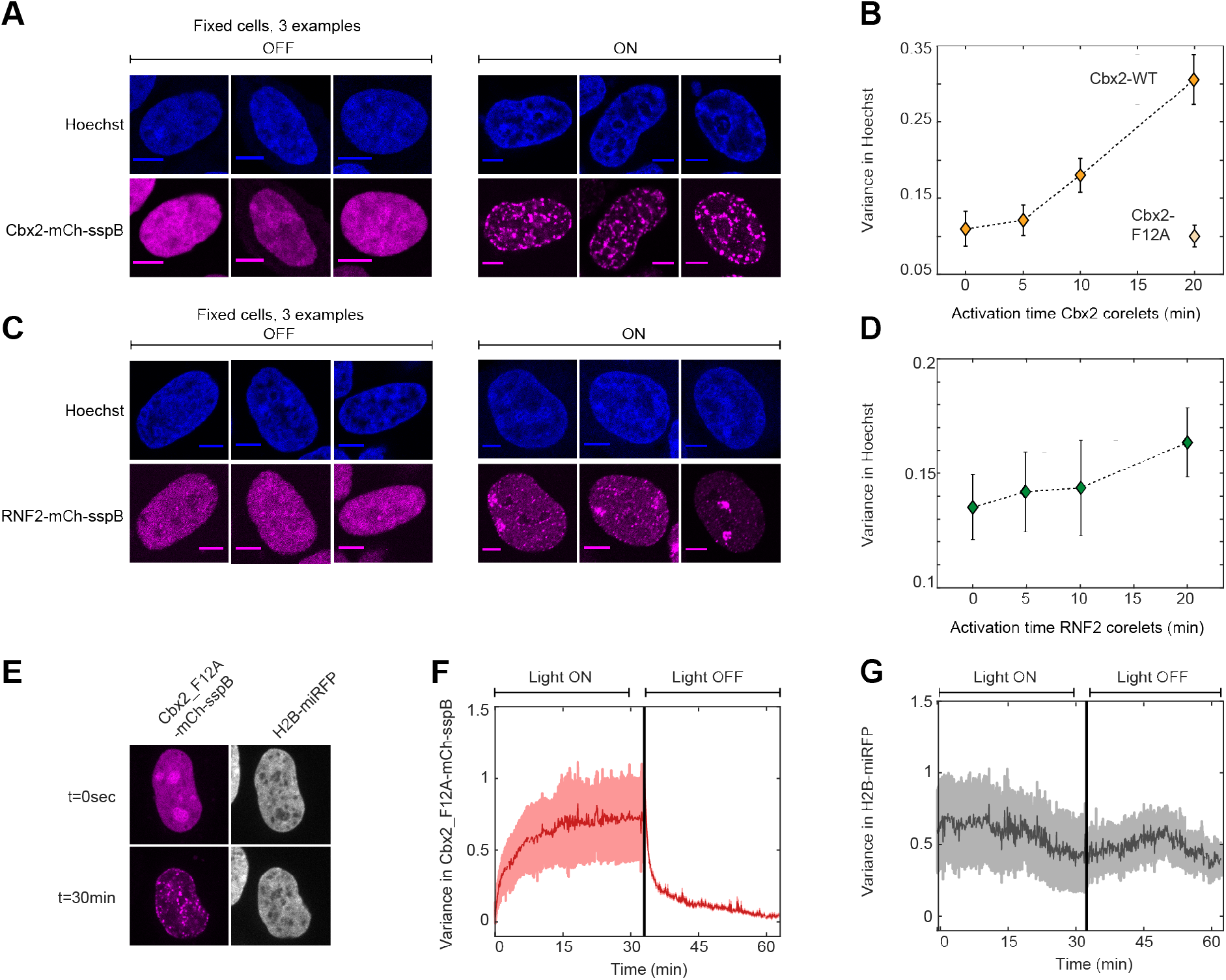
A. Fixed Cbx2-mCh-sspB Corelet cells before (OFF) and after (ON) 20 minutes of activation, stained with Hoechst. Before activation Cbx2-mCh-sspB associates with homogeneously distributed chromatin throughout the nucleus. After light activation, Cbx2-Corelets compact chromatin locally. B. Quantification of the variance in Hoechst increasing with Cbx2-Corelet activation time. The Cbx2-F12A mutant that is unable to interact with H3K27me3 shows compaction compared to the non-activated situation. C. Before activation (OFF) the RNF2-mCh-sspB is homogeneously distributed throughout the nucleus. After light activation, RNF2-Corelets partition onto the inactive X-chromosome and form small de novo puncta throughout the nucleus. There is little change in the Hoechst distribution. D. Quantification of the variance in Hoechst increasing with RNF2-Corelet activation time. E. Cbx2_F12A-Corelet cell with H2B-miRFP before (t=0) and after activation (30 min). F. The variance in the Cbx2_F12A signal increases rapidly as condensates form, and decreases upon deactivation. G. The variance in the H2B-miRFP is unaffected by the condensates.

## METHODS

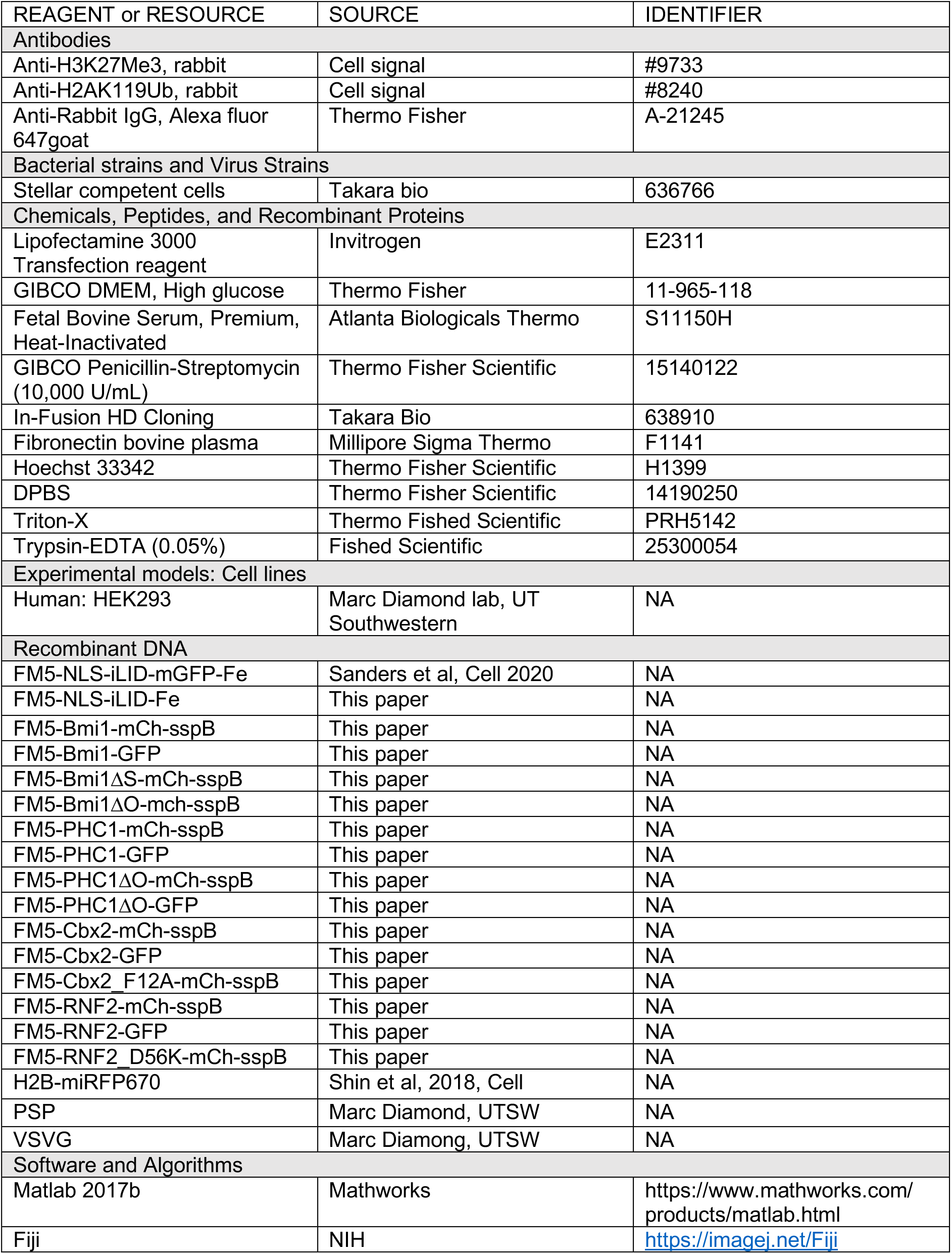

## METHOD DETAILS

### Cell culture

HEK293 cells were used for virus preparation and experiments. Cells were culture in 10% FBS (Atlanta biological) DMEM (GIBCO) supplemented with penicillin and streptomycin at 37°C with 5% CO_2_ in a humidified incubator. On the day before imaging, cultured cells were dissociated with trypsin (Trypsin-EDTA 0.05%, Fisher Scientific) and transferred on a 96-well glass-bottom dish (Thomas Scientific).

### Plasmid construction

DNA fragments encoding our proteins of interest were amplified with PCR using oligonucleotides synthesized by IDT, and CloneAMP HiFi PCR premix. Constructs were cloned using In-Fusion HD cloning kit (Clonetech). Cloning products were confirmed by sequencing.

### Lentiviral transduction

All live cell imaging experiments were performed on stably expressing cells, transduced with lentivirus. Lentiviruses were produced by transfecting the desired construct with helper plasmids VSVG and PSP into HEK293T cells with Lipofectamine 3000 (Invitrogen). Virus was collected 2 days after transfection and used to infect WT HEK293 cells in 96 well plates. Three days after addition of virus, cells were passaged for stable maintenance.

### Live cell imaging

Images were taken on a Nikon A1 laser scanning confocal microscope using a 60x oil objective (Apo 60x/NA 1.4). The microscope was equipped with a stage incubator to keep cells at 37°C and 5% CO_2_. Proteins tagged with Hoechst were imaged using a 405nm laser, GFP with 488nm, mCherry with 560, and Alexa647 with 640nm. Imaging was done on an area of 60×60μm (512/512 pixels).

### Corelet activation (global, local, offline)

Cells were captured in the mCherry channel only, to visualize the sspB component before activation. Cells were then activated with the 488nm laser (typically for 3 minutes) with a frame interval of 2 seconds, while imaging in GFP and mCherry at Nyquist zoom. For local activation, a 1μm^2^ square is activated. For following compaction over time, the 3 minute activation was followed by 30 minutes of 5 second intervals of activation.

For offline activation, cells were placed on an LED array (Amuza). After activation for an indicated amount of time, 4% formaldehyde is added to the wells. After 10 minutes incubation, cells are washed with PBS. The cells are then treated with 1:2000 Hoechst in PBS for 20 minutes, or continued to immunofluorescence.

### Fluorescence recovery after photobleaching

Cells were activated globally as described above to acquire steady state. A 1μm^2^ area in the cluster was then bleached with the 560nm laser to quench the mCh-sspB component of the condensate. Fluorescence recovery was followed while imaging in both mCh and GFP channels. FRAP experiments were analysed by measuring the mean fluorescence intensity in the bleached area (1μm^2^) over time. Error bars represent standard deviation on at least 5 replicates.

### Immunofluorescence

After activation and fixation as described above, cells were washed with washbuffer (0.35% Triton-X in PBS) for 5 minutes, and permeabilized with 0.5% Triton-X in PBS for 1 hour.

Cells were then blocked for 1 hour using blocking buffer (0.25% Triton-X, 5% FBS, in PBS). Primary antibodies were dissolved in blocking buffer (all 1:1000) and incubated overnight at 4°C. The next day, cells were washed 3x for 5 minutes with washing buffer. The secondary antibody (AlexaFluor 647 goat-anti rabbit, 1:1000) was dissolved in blocking buffer, and incubated for 2 hours at 4°C. Cells were washed 3x for 5 minutes with washbuffer, followed by 20 minutes incubation of 1:2000 Hoechst. Finally, cells were washed with PBS.

### RNA FISH

The Stellaris RNA FISH protocol for Adherent Cells was used for detecting XIST in the cells with RNF2-GFP. The growth medium was decanted, the cells were washed with PBS and then fixed using 4% formaldehyde. After incubation for 10 minutes at room temperature, the cells were washed twice with PBS and then permeabilized by adding 70% ethanol for a minimum of an hour.

For hybridization, the ethanol was decanted and cells were washed according to manufacturer’s protocol, and left to incubate at room temperature for 5 minutes. Washbuffer was then decanted and the Hybridization Buffer (10% formamide in Stellaris RNA FISH Hybridization Buffer) containing 2% of the Stellaris RNA FISH Probe (Human XIST probe with Quasar 570 Dye) re-dissolved in TE Buffer (10 mM Tris-HCL, 1 mM EDTA, pH 8.0) was added and incubated in the dark at 37°C for 16 hours. The Hybridization Buffer was then aspirated, Washbuffer A was added and the cells were left to incubate in the dark at 37°C for 30 minutes. Finally, the buffer was decanted and cells were imaged as described.

### Image analysis

All images were analysed using a combination of manual segmentation (ImageJ) and automated segmentation in Matlab.

### Phase-diagram construction

A standard imaging protocol was used on all cells to avoid variability. All activation protocols were 3 minutes with 2 second intervals. Only cells that were fully in the field of view were considered. The average GFP and mCh fluorescence intensity was determined using the first frame (before activation). All measurements were confirmed by an independent researcher. FCS calibration curves were used to determine the mCh and GFP concentrations from the fluorescence intensities as described in Bracha et al (2018).

Valence was measured as the ratio of sspB-fused protein to core.

### Pearson-correlation coefficient and variance determination

To estimate the degree of colocalization between two channels, first, the nucleus was segmented with a custom Matlab script. As they can heavily skew the degree of colocalization, the nucleoli and inactivated X-chromosome were excluded. Then, the pixel-by-pixel correlation was determined using the Pearson-correlation coefficient (PCC) as defined as:

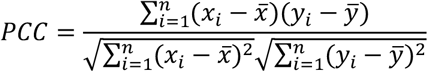

Where *n* is the sample size (number of pixels), and *i* is the individual pixel index, with *x*_*i*_ and *y*_*i*_ the value in that pixel for the two channels, and 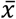 and 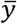 the sample means.

In the case of co-expressed PRC proteins (shown in Figure 2), the analysis was done in a similar manner, with the exception that the inactivated X-chromosome was not excluded from analysis. The symmetry of the interactions can be seen from the fact that all data points lie relatively close to the diagonal.

For variance determination, the nucleus was segmented in every frame, with nucleoli excluded. The variance was determined in each channel.

